# Theta phase specific modulation of hippocampal memory neurons

**DOI:** 10.1101/2022.10.27.513992

**Authors:** Bahar Rahsepar, Jad Noueihed, Jacob F. Norman, Benjamin Lahner, Melanie H. Quick, Kevin Ghaemi, Aashna Pandya, Fernando R. Fernandez, Steve Ramirez, John A. White

## Abstract

The theta rhythm, a quasi-periodic 4-10 Hz oscillation, is observed during memory processing in the hippocampus, with different phases of theta hypothesized to separate independent streams of information related to the encoding and recall of memories. At the cellular level, the discovery of hippocampal memory cells (engram neurons), as well as the modulation of memory recall through optogenetic activation of these cells, has provided evidence that certain memories are stored, in part, in a sparse ensemble of neurons in the hippocampus. In previous research, however, engram reactivation has been carried out using open loop stimulation at fixed frequencies; the relationship between engram neuron reactivation and ongoing network oscillations has not been taken into consideration. To address this concern, we implemented a closed-loop reactivation of engram neurons that enabled phase-specific stimulation relative to theta oscillations in the local field potential. Using this real-time approach, we tested the impact of activating engram neurons during the peak (encoding phase) and trough (recall phase) of theta oscillations. Consistent with previously hypothesized functions of theta oscillations in memory function, we show that stimulating engram neurons at the trough of theta is more effective in eliciting behavioral recall than either fixed frequency stimulation or stimulation at the peak of theta. Moreover, phase-specific trough stimulation is accompanied by an increase in the coupling between gamma and theta oscillations in CA1 hippocampus. Oure results provide a causal link between phase-specific activation of engram cells and the behavioral expression of memory.

## Introduction

The discovery of neurons whose activity correlates with memory activity, often referred to as engram cells, has offered evidence that certain memories are stored in a sparse set of neuronal ensembles across the hippocampus (Reijmers et al. 2007). These cells are active during the encoding of a memory and reactivate upon recall of that specific event (Denny et al. 2014; Tayler et al. 2013). Further, artificial reactivation of engram neurons induces recall-like behavior, thus establishing a causal role for these neurons in memory processing (Liu et al. 2012; Ramirez et al. 2013).

Previous studies have utilized activity based neural tagging strategies, which link expression of a protein of interest (e.g. Channelrhodopsin) to the expression of immediate early genes (IEG) (most commonly Arc (Denny et al. 2014) and cFos (Liu et al. 2012)) to tag and modulate active populations during a specific event. The modulation of memories through activity based tagging of engram neurons has often targeted cells in the hippocampus (Josselyn and Tonegawa 2020; Tonegawa et al. 2015); a brain structure in the temporal lobe with a modular design that consists of three major sub-regions: the dentate gyrus (DG), CA3, and CA1, all with extensive interconnectivity (Andersen et al. 2009; LR, CE, and RE 2004). Each of the sub-regions has distinct connectivity patterns that potentially provide unique stages in the processing of memories.

Recently, studies have probed this functional organization by driving engram neurons in different regions of the hippocampus, such as DG (Liu et al. 2012; Ramirez et al. 2013) and CA1 (Redondo et al. 2014; Ryan et al. 2015). In these studies, however, there have been discrepancies in the most effective stimulation frequency, both between regions as well as within regions (Compare Ohkawa et al., 2015; Ryan et al., 2015). Further, studies have been limited to activating engram neurons using fixed frequency stimulation, without taking into consideration the ongoing spontaneous network activity. In particular, hippocampal regions are dominated by a quasi-periodic 4-10 Hz network-wide theta oscillation in the field potential generated by temporally organized firing during memory processing (Buzsáki 2002; Colgin 2013).

The hippocampus is tasked with both encoding of new information, as well as recalling past experiences. A prominent model, termed the Separate Phase of Encoding and Recall (SPEAR) model (Hasselmo, Bodeí, and Wyble 2002), posits that encoding and retrieval are temporally interleaved at different phases of hippocampal theta oscillations. In the SPEAR model (Hasselmo et al. 2002; Hasselmo and Stern 2014), the peak of the hippocampal theta oscillation (as measured in *striatum pyramidale*) is dominated by inputs from the entorhinal cortex, which carry sensory information that is potentially required for the encoding process. In contrast, the trough (negative peak) of theta oscillations occurs during strong CA3 activity, a region of the hippocampus known for pattern completion that is supported through strong recurrent connections (Leutgeb et al. 2007; Rolls 2016; Senzai 2019), and thus ideally suited for the retrieval of previously stored memories.

The SPEAR model has been supported by electrophysiological data *in vivo* (Hyman et al. 2003) and *in vitro* (Kwag and Paulsen 2009). These studies have indicated that the peak of theta oscillations is associated with strong long-term potentiation, which can support the encoding process, while the trough of theta has strong long-term depression that supports the recall of previously stored memories (Douchamps et al. 2013; Manns et al. 2007). In behaving mice, memory performance can be altered through phase-specific inhibition of neurons in CA1, the output region of the hippocampus (Siegle and Wilson 2014). Finally, a recent study in human subjects has shown a strong correlation between memory tasks and the phase of theta (Kragel et al. 2020). Despite these results, as well as the prior development of closed loop optogenetics stimulation (Grosenick, Marshel, and Deisseroth 2015), the activation of engram neurons has been carried out at fixed frequencies, without taking into account the phase of theta oscillations.

Here, we address this problem by tying the modulation of engram neurons to specific phases of theta oscillations. We implemented a real-time phase prediction algorithm and tested the behavioral and electrophysiological effects of theta phase-specific stimulation in gating memory recall. We compared results from phase-specific stimulation with those gathered using fixed-frequency stimulation at the previously established value of 20 Hz, as well as stimulation at 6 Hz, which provided a control representing the average stimulation frequency during phase-specific activation. Our results support the SPEAR model, with optogenetic stimulation of engram neurons in DG at the trough (recall) phase driving the most robust behavioral recall and the largest amount of coupling between the theta and gamma rhythms in CA1 region.

## II. Results

### 1. Closed loop photo stimulation at specific phases of theta oscillations

To date, engram work has used fixed frequency optogenetic stimulation to reactivate tagged cells and drive behavior linked to memory recall. For example, most studies have used 20 Hz stimulation, which is well above the natural firing rates measured in the dentate gyrus (1 Hz on average, with peak rates of 8 Hz) (GoodSmith et al. 2017; Senzai and Buzsáki 2017). We hypothesized that a more physiologically realistic stimulus pattern, both in terms of firing rate and timing, would reactivate memories more effectivity. To test our hypothesis, we developed a protocol that phase-locked the stimulation time to the theta rhythm.

To deliver photo-stimulation at specific phases of the theta oscillation, we used a custom a real-time phase detection-and-prediction algorithm. Our algorithm (Figure 1) reads the LFP from the CA1 region of the hippocampus and filters it using a finite impulse response filter between 4 and 10 Hz implemented in the Real-Time eXperimental Interface (RTXI) software (Lin et al. 2010). The algorithm predicts the timing of the next desired extremum by averaging the duration of previous cycles and sends a TTL pulse to drive the laser at the predicted time of either the peak or the trough of the next theta cycle. The phase of theta oscillations is in reference to measures in *striatum pyramidale* in the hippocampus. If the location of the electrode, as determined by post-hoc histological analysis (see methods), was observed to be in another layer of the hippocampus, we corrected for this change (trough vs. peak) to ensure similar theta phase across animals.

**Figure 1.**
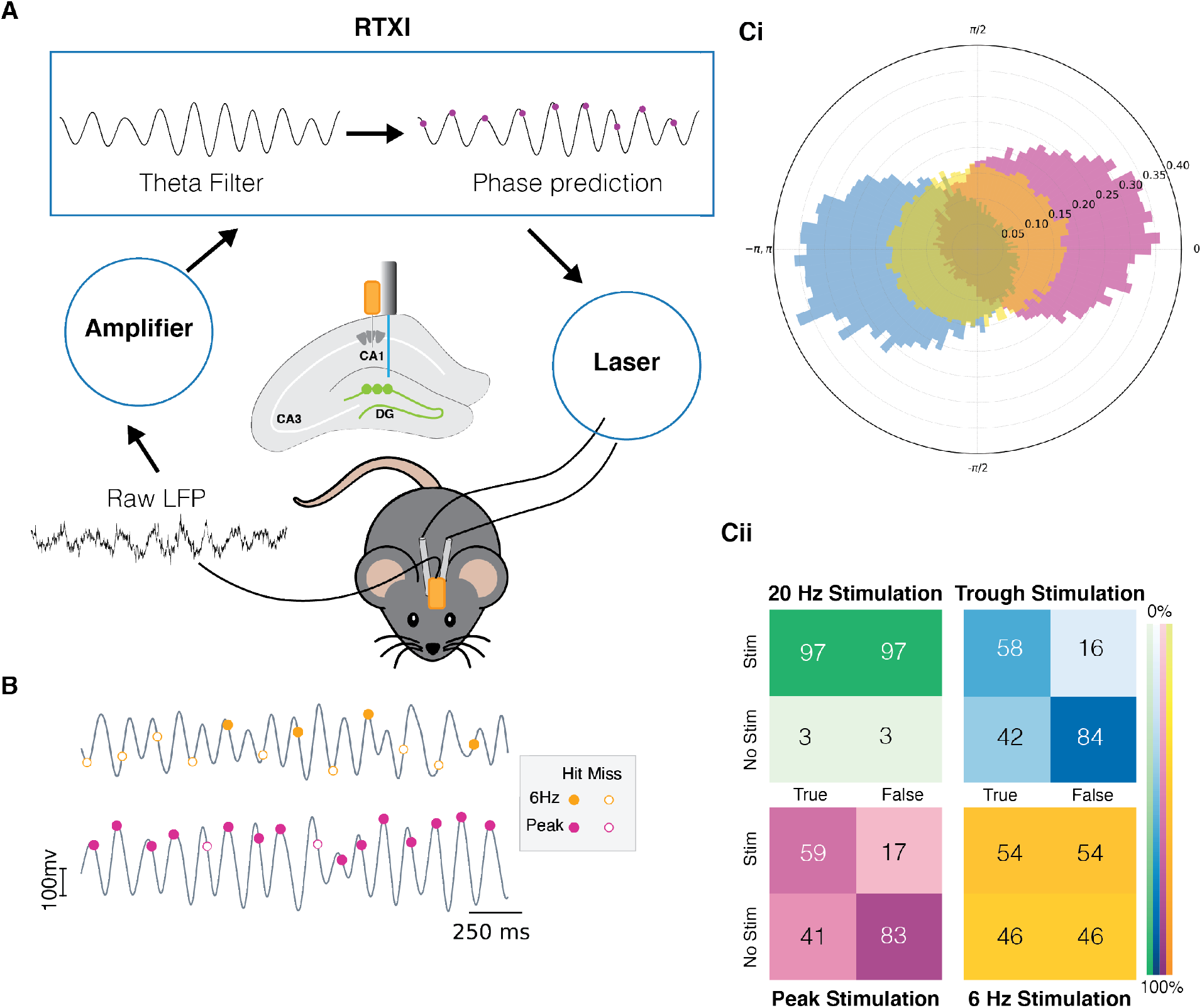
Quantification of the real-time phase detection algorithm performance. ***A***. LFP signal recorded from the hippocampus is amplified and processed in Real-Time eXperimental Interface (RTXI). The signal is first filtered using a FIR filter in the theta range (4-12 Hz) and then a real-time phase detection algorithm predicts the next extrema. At the predicted time, a TTL pulse is sent to the laser which delivers light through fiber optics to the DG region of the hippocampus to activate tagged neurons. ***B***. Sample stimulation shows the superior performance of the predictive algorithm in comparison to the 6 Hz periodic stimulation. ***C. (Ci)*** Normalized polar histogram shows the phases of stimulation in the cases of peak, trough, and periodic 20 Hz and 6 Hz stimulation. ***(Cii)*** Confusion matrices indicate that peak and trough stimulation are specific to the desired phase of the stimulation.

The average frequency during both peak- and trough-specific stimulation was 6 Hz. Hence, we compared the performance of our real-time algorithm and phase-specific stimulation with fixed-frequency stimulation at 6 Hz. We also performed experiments with 20 Hz stimulation, as this is a commonly used stimulation frequency in past studies. As shown in Figure 1, the use of closed loop stimulation resulted in significantly more phase specific stimulation: 83, 84% True Negative Rate (TNR) for peak and trough stimulation, respectively, as compared to 46% and 3% TNR with 6 Hz and 20 Hz stimulations, respectively. These results indicate that our algorithm has a low rate of stimulation outside of the desired phase, which is critical to testing our hypothesis. Moreover, sensitivity of the algorithm is moderately superior to the 6 Hz stimulation with a True Positive Rate (TPR) of 58% and 59% for peak and trough, respectively, as compared to 54% for 6 Hz. Since a 20 Hz stimulation rate is about two times higher frequency than theta oscillations, it results in a constant, non-specific stimulation (TNR and TPR of 3% and 97%, respectively). The specificity of the algorithm is critically important in ensuring we are not confounding the results with a high rate of stimulation at the opposite phase. In terms of sensitivity, a higher than 50 % value from the algorithm is satisfactory as we do not expect that during natural memory processing the animal is encoding or recalling the memory during every theta cycle. Rather, when either of these processes happen, they are *preferentially* happening at either the peak or trough, as explained above.

### 2. Experimental setup for comparing different modes of stimulation

We developed an experimental setup to compare phase-specific and fixed-frequency reactivation of hippocampal engram neurons. Importantly, our setup was designed to test different modes of stimulation within the same animal. Further, as repeated engram reactivation can affect the original memory, our experimental design randomized the order of the stimulation in different animals. The experiment took place in two different contexts: a neutral context A and a fearful context B. As detailed in the Methods section, the two different contexts were differentiated based on a variety of sensory stimuli (see Methods). Animals were first habituated to handling and exploration in the neutral context A. Habituation took place over 4 days, during which both fiber optics and the LFP electrodes were attached to acclimatize the mice to the setup. As illustrated in Figure 2A, the animals were anxious on day 1 as shown by the higher amount of freezing (15%). However, they quickly became habituated to the setup as shown by a much-reduced level of freezing on the last two days of the habituation (5%).

**Figure 2.**
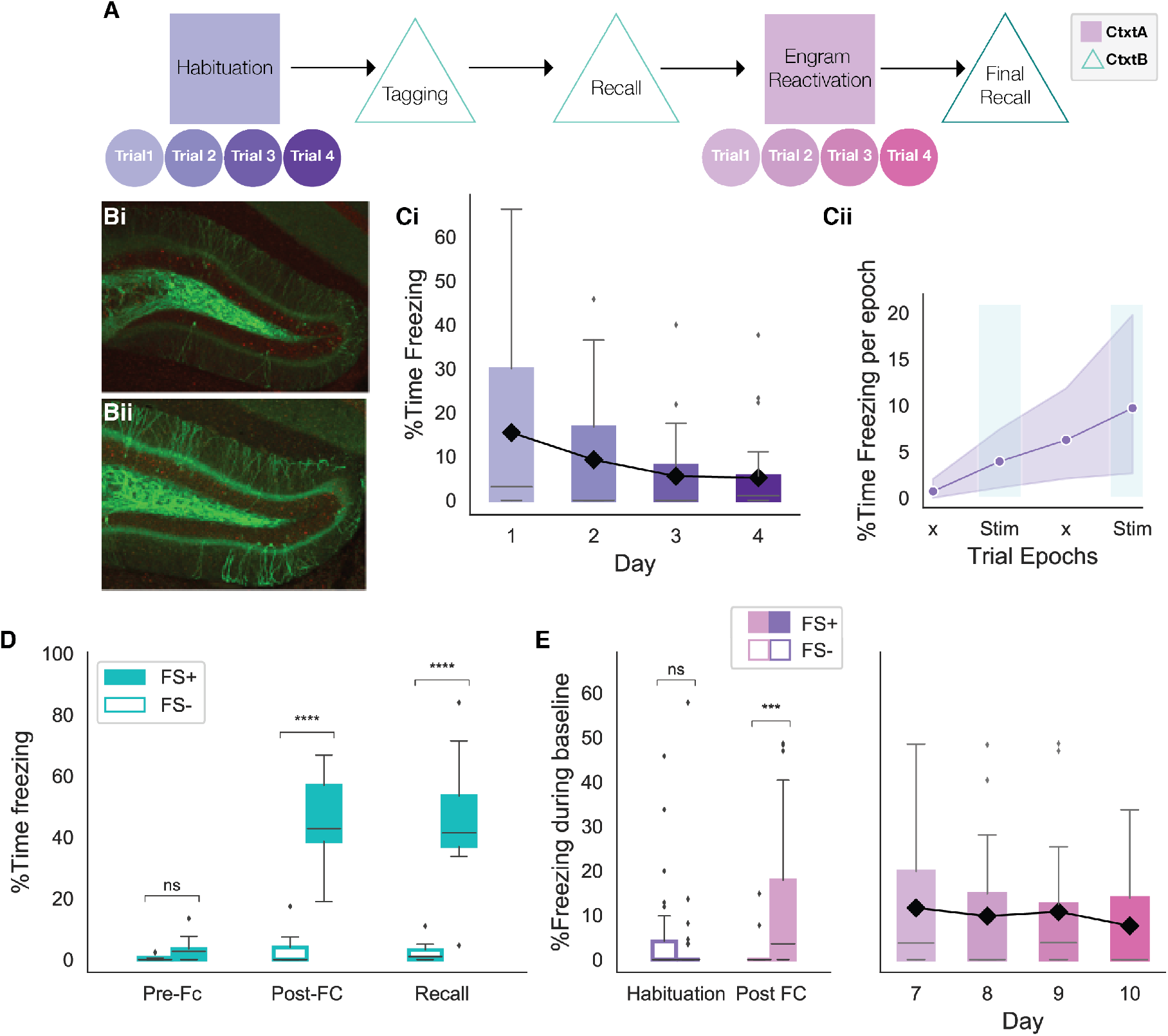
Behavioral experiment design. ***A***. Schematic of the behavioral experiment. Animals were habituated in context A for 4 days prior to the tagging of engram neurons in context B. Following recall, animals were re-exposed to context A and underwent reactivation of the engram neurons with 4 different stimulation strategies on distinct days (trials). The experiment concluded with a final recall in context B. ***B***. Experimental ***(Bi)*** and control ***(Bii)*** animals had a similar number of DG cells tagged as shown by the number of GCs expressing EYFP-ChR2. Experimental animals received a foot shock in context A, while control animals did not. ***C***. Habituation of animals over four days in context A resulted in decreased in freezing, indicating comfort with the setup. Black line shows the trend for the mean. ***Cii***. Average of the percent time freezing over 4 epochs of trials on the last day of habituation (day 4) indicate 5% increase in percent time freezing due to fatigue later in the trial. This value serves as a baseline for subsequent analysis. ***D***. Both experimental (FS+, n = 17) and control (FS-, n = 9) animals showed minimal freezing in context B prior to the foot shocks (independent t-test, p = 0.7). However, experimental animals showed significantly higher freezing post foot shock that persisted on the following day, indicating recall (****p < 0.00001). ***E***. On the last day of habituation, both FS+ and FS-groups exhibited minimal baseline freezing (p = 0.6). However, post fear conditioning, the FS+ group showed significantly higher baseline freezing (***p < 0.0001). The elevated baseline freezing for the experimental animals is sustained throughout all 4 days of the experiment. The black line shows the trend for the mean freezing. (For all figures, box shows the quartiles of the dataset, while whiskers show the rest of the distribution. Outliers are shown using diamonds)

Each trial of the habituation was designed to be the same length as the experimental trials, consisting of four 3-minute epochs in which the stimulus lights were turned off and on. Animal freezing on the last day of habituation was used as a measure of baseline freezing change for the rest of the experiment. As shown in Figure 2B, freezing increased by about 5% during the trial. Following habituation, animals were taken off doxycycline 48 hours prior to fear conditioning to allow for the tagging of engram neurons. On the tagging day, animals freely explored context B for 5 minutes (Figure 2D, pre-FC). Over a second, 5-minute interval, experimental animals (FS+) received four 1.2 mA foot shocks, while the control animals (FS-) were left to explore freely. Both experimental and control animals showed minimal freezing prior to the shocks, with only experimental animals exhibiting elevated freezing after the foot shocks (Figure 2D). Post fear conditioning, animals were put back on a doxycycline diet and returned to a new home cage. On the following day, animal recall of the fearful context was tested by re-introducing them to the fearful context B for 5 minutes. As shown in Figure 2D, only FS+ animals show an elevated level of freezing, indicating successful recall of the fearful memory.

Following tagging and recall of the fearful memory in context B, animals were re-exposed to the neutral context A in which they were originally habituated. Post-fear conditioning, only experimental animals showed elevated baseline freezing (1.5% pre vs 10% post), indicating a potential generalization of the fearful context B memory to the neutral context A (Figure 2E). Despite the elevated freezing (10%), neutral context A freezing was lower than the freezing following fear conditioning (45%) or during recall (46%) of the fearful memory in context B (Figure 2D vs. 2E). However, the increased baseline freezing in neutral context A obscured the light induced freezing and, therefore, required the artificial memory reactivation to generate higher freezing levels to be deemed effective. It is important to note that the elevated baseline freezing is similar across days, making it possible to pool data from different days. As a result of the elevated baseline freezing, an effective stimulation needed to be powerful enough to elicit a behavioral response (increase in percent time freezing) beyond the elevated baseline.

### 3. Trough stimulation leads to stronger and more robust recall

Next, we compared the effects of engram reactivation via different modes of stimulation using phase-specific and fixed frequency stimulation. Because we did not detect any differences across measures taken at different days, we pooled the data across days. As shown in Figure 3Ai, only stimulation at the trough of theta could drive the expected increase in freezing during both stimulation epochs. Although 6 Hz stimulation was effective during the first stimulation epoch, this frequency failed to elicit significant freezing during the second epoch. Peak and 20 Hz stimulations showed a gradual increase in freezing that was similar to the habituation trial. Averaging the light induced freezing across epochs indicated that only trough stimulation resulted in significantly higher freezing rates (Figure 3Aii; paired t-test, p < 0.01).

**Figure 3.**
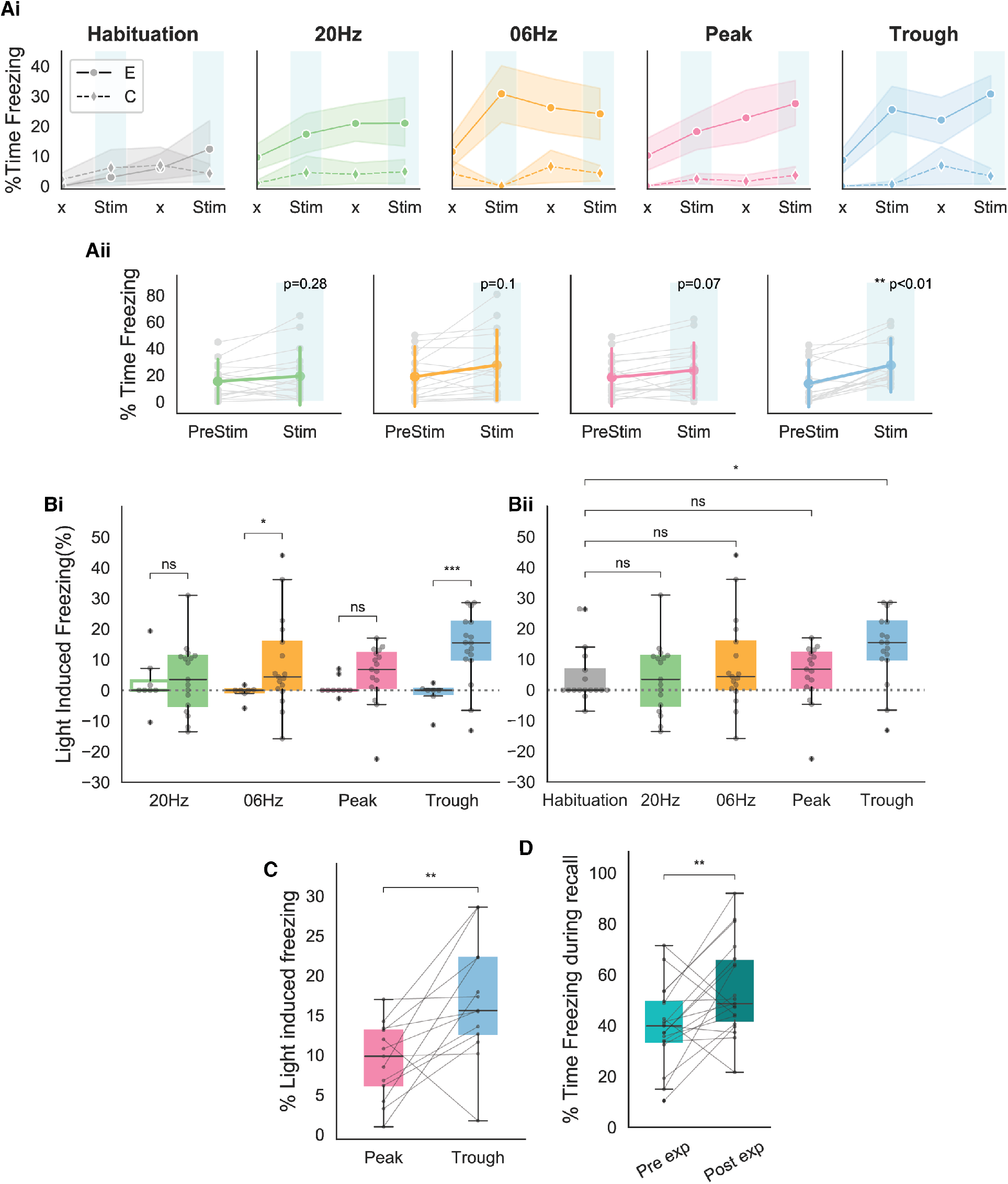
Behavioral responses indicate that recall is largest when using stimulating at the trough of theta. ***A. (Ai)*** Average freezing per epoch for FS+ (solid line) and FS-(dashed line) animals during habituation (grey) and during the four modes of stimulation. (20 Hz: Green, 6Hz: Yellow, Peak: Pink, Trough: Blue). ***(Aii)*** Average increase in freezing using no stimulation (epochs 1 and 3) and stimulation (epochs 2 and 4). Paired t-test indicates that only trough stimulation reliably caused increased freezing that resulted from activation of engram neurons. ***B. (Bi)*** Average light induced freezing was calculated for both experimental (FS+) and control (FS-) animals by subtracting epochs 2 and 4 from epochs 1 and 3, respectively, and averaging the value. Only 6 Hz stimulation (independent t-test, * p = 0.02 < 0.05) and trough stimulation (***p=0.0002 < 0.0001) showed light induced freezing that differed significantly from the non-foot shocked group. Light induced freezing of using peak and 20 Hz stimulation was not significantly different than the control group (20Hz: p = 0.6, Peak: p = 0.07). ***(Bii)*** Light induced freezing on the last day of habituation prior to the experiment only differed significantly for trough stimulation. (20 Hz: p = 0.8, 6 Hz: p = 0.4, Peak: p = 0.8, Trough: *p=0.02 < 0.05). ***C***. Paired comparison between trough and peak stimulation for animals that exhibited light induced freezing indicated significantly higher levels of freezing induced by trough stimulation (n = 13, **p=0.007 < 0.001). ***D***. Significantly higher freezing was observed upon exposure to the fearful context B 4 days after artificial reactivation of engram neurons in context A (paired t-test, **p < 0.01).

To compare the effects of the stimulation between different stimulation patterns in the control and experimental animals, as well as between different stimulation patterns, we calculated the percent of light induced freezing. For this measure, the change in freezing relative to the baseline prior to the stimulation was calculated by subtracting the percent time freezing in epochs 2 and 4 from epochs 1 and 3, respectively. The average of these two values is referred to as light induced freezing. As shown in Figure 3B, only 6 Hz and trough stimulations resulted in significantly higher values of light-induced freezing when compared with the control animals receiving the same stimulation. Further, paired comparisons of the amount of light induced freezing during each stimulation indicated that only trough stimulation elicits light induced freezing values significantly different than the habituation trials. As a result, the increase in freezing in the cases of peak stimulation, as well as 20 and 6 Hz fixed frequency stimulations, were like those expected from a general increase in the animal’s immobility in later trials (Figure 2C).

Direct comparisons of trough and peak stimulations within each animal showed significantly higher freezing values when using trough stimulation (Figure 3C). For this analysis, we focused only on the cases in which both the peak and trough caused light-induced freezing. The analysis indicated that activating engram neurons at the trough of theta was more effective at inducing artificial recall of a tagged memory. We also found that the experimental group showed significantly higher freezing post reactivation of engrams in the neutral context A, which we believe indicates a potential strengthening of the original tagged memory.

### 4. Electrophysiological hallmarks during different forms stimulation indicate changes in gamma-theta coupling following trough stimulation

Having established that trough stimulation is the most effective stimulus in driving freezing behavior, we sought to identify physiological hallmarks of its efficacy. To start, we analyzed the LFP recordings from the CA1 region of the hippocampus. Consistent with previous studies, theta oscillations were higher in frequency and larger in amplitude during locomotion (Figure 4B, see the ‘bumps” in the blue lines in the right panels) when compared with measurements made during freezing (red lines). This trend held during all epochs of the experiment regardless of the presence or absence of the stimulation. Similarly, results were not different when using different modes of stimulation (data not shown).

**Figure 4.**
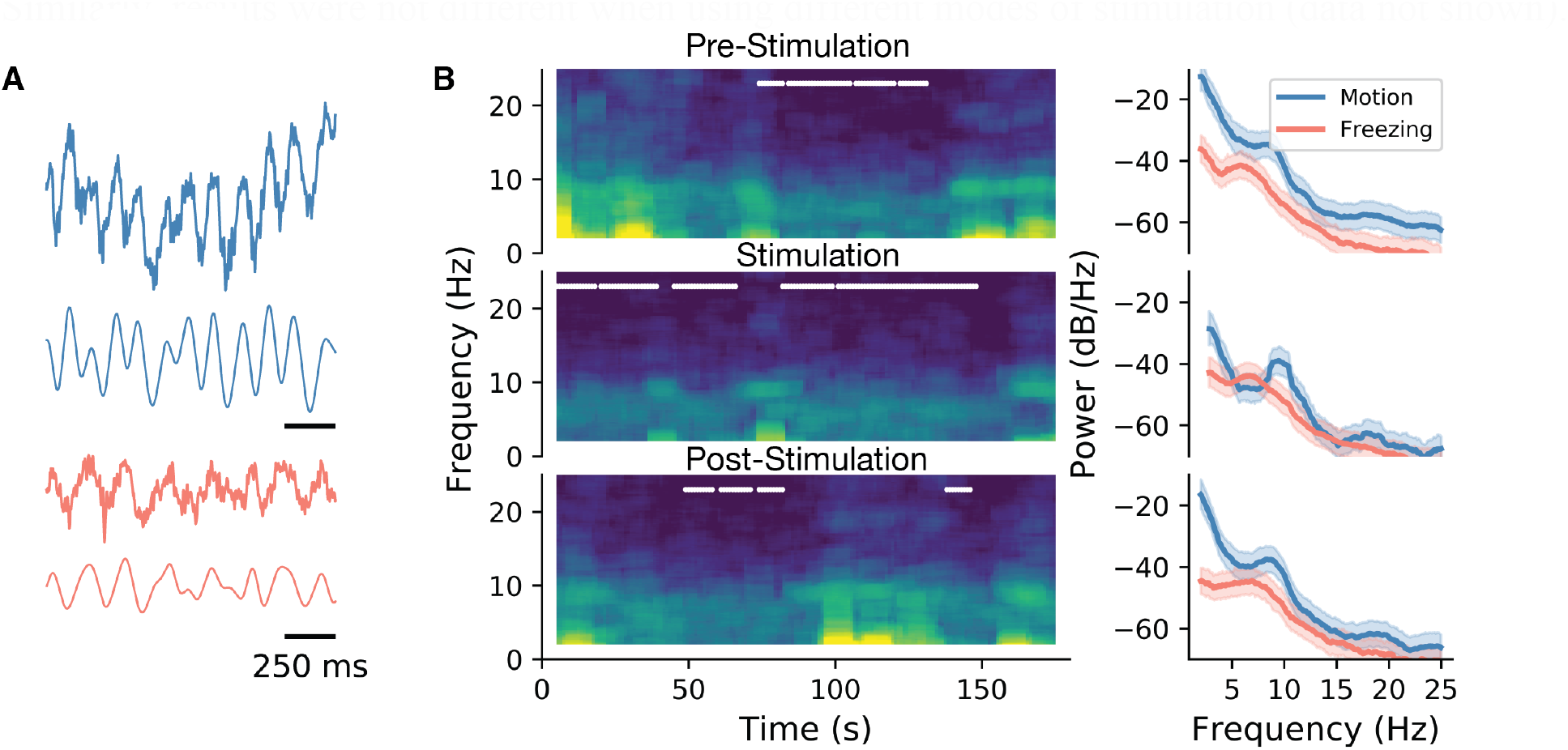
LFP characteristics during locomotion and stimulation. ***A***. Sample LFP recordings from CA1 during active exploration (blue) and freezing (red). ***B***. Spectrogram for the whole duration of an epoch (left), and the associated power spectral density graph. In spectrograms, freezing episodes are marked with a white line. As shown, during all three epochs (pre-stimulation: epoch 1, stimulation: epoch 2 and post-stimulation: epoch 3), power and the peak frequency of theta were lower during freezing episodes. This is indicated more clearly in the PSD graphs on the right as evidenced by the shifts in the peak of theta oscillations.

The 30 - 100 Hz gamma rhythm has been theorized to play a critical role in hippocampal memory processing (Lisman and Jensen 2013). Electrophysiological studies have established that theta-gamma coherence is correlated with coordinated information transfer between different sub-regions of the hippocampus (Pernía-Andrade and Jonas 2014). Moreover, cross-frequency coupling (CFC) has been demonstrated in a memory test experiment that established a correlation between the strength of this coupling and memory performance (Tort et al. 2009). These studies quantified the CFC using a metric termed the Modulation Index (MI) (Tort et al. 2010), which is calculated by measuring the distribution of the gamma amplitude within specific phases of theta. This is a measure of Phase Amplitude Coupling (PAC) between theta and gamma oscillation. To test the role of gamma oscillations, we applied the MI metric to our dataset and tracked its correlation to the efficacy of artificial memory modulation eliciting recall.

Our analysis indicated that the MI was highest between the phase of theta filtered at 4-8 Hz frequency and the amplitude of 55-85 Hz gamma, known as mid gamma. This is similar to previous observations in CA1 (Jiang et al. 2020; Schomburg et al. 2014; Zhang et al. 2019) and consistent with CA3 inputs to CA1 driving recall of memories (Colgin 2015, 2016). After establishing the presence of this cross-frequency coupling, we sought to quantify its value during different epochs of the experiment, as well as during different modes of engram reactivation. Comparing the modulation index at baseline (pre-stimulation epoch) indicated no difference between the four stimulation patterns. However, during the stimulation period, only trough activation showed elevated values in MI, which then went back to control levels during the post-stimulation period (epoch 3, Figure 5B). Comparing pre-stimulation to stimulation epochs for each mouse confirmed that the MI was only significantly modulated in cases using trough stimulation, which was significantly higher than both pre and post stimulation. In other stimulation setups, we observed no differences in the MI between the three epochs (Figure 5C).

**Figure 5.**
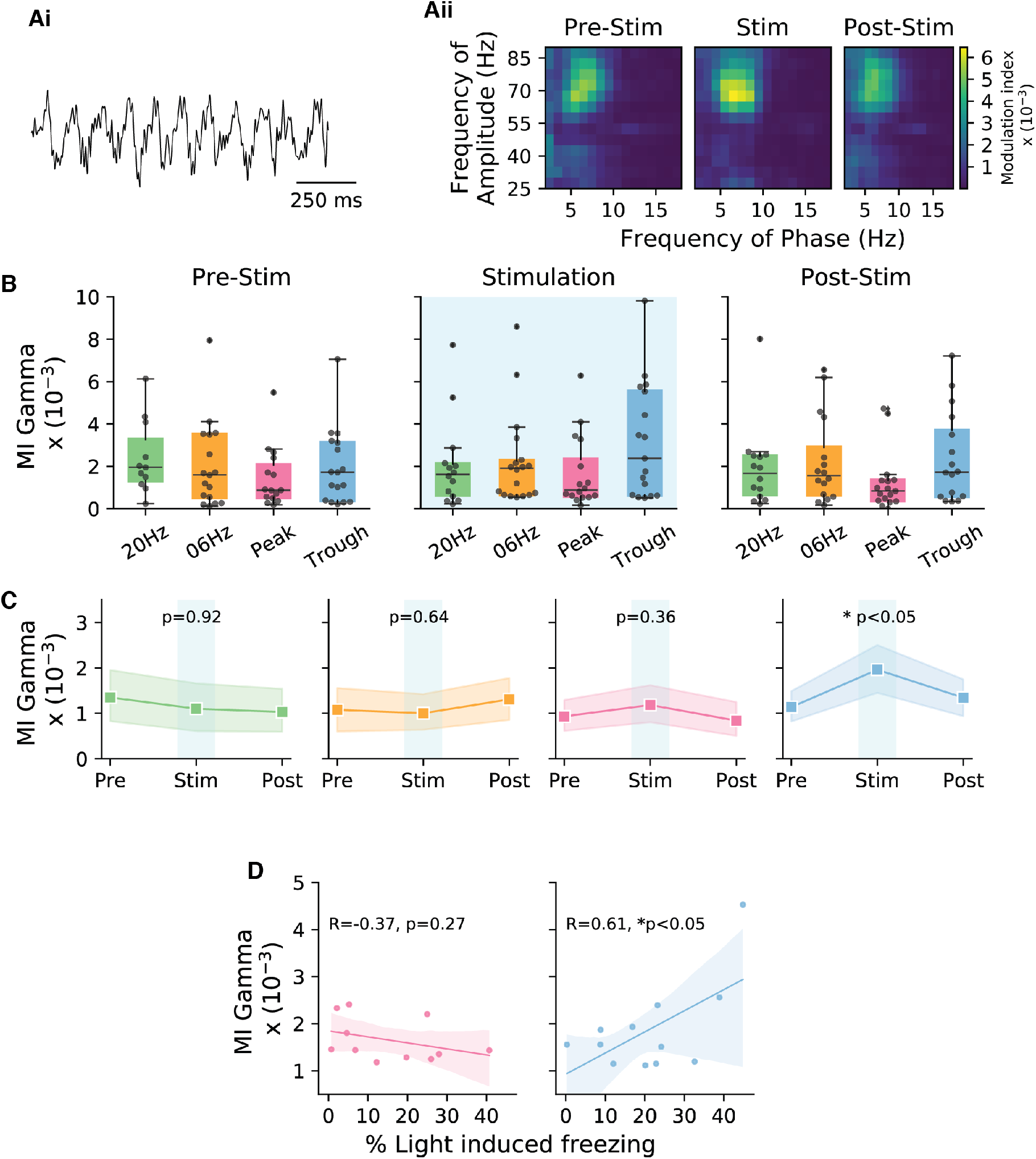
Theta-Gamma cross frequency coupling correlates with memory recall performance during trough stimulation. ***A. Ai***. Sample LFP recording indicating theta-nested gamma oscillations. ***Aii***. The modulation index was calculated for cross-frequency coupling between the gamma amplitude and the phase of theta for trough stimulation during epochs 1-3 corresponding to pre-stimulation, stimulation, and post-stimulation. Comodulograms only showed an increase in the MI during the stimulation epoch. ***B***. Boxplots show the modulation index during the three different epochs for the four stimulation modes. (n = 17). ***C***. Direct comparison of the three epochs for the different stimulation modes indicates that only trough stimulation significantly increased the MI during the stimulation epoch. ***D***. Correlation of the stimulation efficacy (% light induced freezing) and MI was only correlated significantly in the case of trough stimulation, and not during peak stimulation. Note, only mice that showed light-induced freezing were included in the analysis (Peak: n = 11, Trough: n = 12).

Consistent with the SPEAR model, our data support the hypothesis that the peak and trough of theta correspond to different modes of hippocampal function with regards to memory processing. In particular, we observed that with trough stimulation the behavioral response was stronger (higher light induced freezing) and that the MI was higher (Figure 5D, left). This relationship was in the opposite direction, albeit not at a significant level, when using stimulation at the peak of theta (Figure 5D, right).

## III. Discussion

Using engram neuron tagging and theta phase specific photo-stimulation, we assessed the functional role of theta phase in memory processing. We show that activating engram cells during the trough of the theta field potential is most effective at inducing recall of stored memories and yields the most robust behavioral outcome corresponding to successful artificial reactivation of the tagged memory. When artificial recall is elicited through phase-specific stimulation, the behavioral outcome is well correlated with an increase in phase-amplitude coupling between theta and gamma oscillations, which has been established as an electrophysiological correlate of memory performance (Kragel et al. 2020; Tort et al. 2009).

Previous studies have observed a frequency-dependency in eliciting behavioral responses across different regions of the hippocampus. For example, Ryan et al. 2015 activated CA1 engram neurons using 4 Hz optogenetic stimulation and observed consistent behavioral responses, while Ohkawa et al. 2015 were able to drive behavior with 20 Hz stimulation in the same region. In the case of DG, the majority of past studies have used 20 Hz stimulation to drive engram activation despite the rate being much higher than natural DG granule cell firing rates in vivo (GoodSmith et al. 2017; Senzai and Buzsáki 2017).

In our current work, we quantified the effect of stimulation frequency on recall by comparing closed- and open-loop approaches of engram neuron reactivation that generated different average stimulation frequencies. Although 6 Hz stimulation in the open loop configuration was moderately effective in eliciting recall, it only occurred when the baseline freezing rate was low. It should be noted that this mode of activation is close to the peak physiological firing rates of the DG granule and mossy cells (GoodSmith et al. 2017; Senzai and Buzsáki 2017). Despite the success of 20 Hz stimulation in artificial recall of the tagged memory in DG (Liu et al. 2012; Ohkawa et al. 2015; Ramirez et al. 2013; Redondo et al. 2014), we did not observe significant light induced freezing with this frequency of stimulation. A possible explanation for this discrepancy was the presence of an elevated baseline freezing rate post fear conditioning in the neutral context in our study (average 10% (Figure 2) versus less than 5% in previous studies (e.g. (Cowansage et al. 2014))). As a result, the elevated baseline freezing could be masking the effect of the 20 Hz stimulation. Moreover, based on the distribution of enhanced and inhibited freezing with 20 Hz stimulation (Figure 3), we hypothesize that this frequency of stimulation could both activate or inhibit engram neurons. In support, work measuring the impact of mossy fiber stimulation on CA3 neuron firing rates (Lee et al. 2019) indicates that 20 Hz stimulation of DG mossy fibers can potentially lead to inhibition of CA3 neurons through feedforward inhibition. For this reason, the potential inhibition caused by 20 Hz stimulation could lead to a lack of stimulation efficacy. In contrast, Lee et al. also showed that 6 Hz stimulation can have a net positive effect on CA3 neuron activity, which is consistent with us observing elevated light induced freezing during 6 Hz stimulation (Lee et al. 2019). Testing all four different forms of stimulation within a single subject to control for inter-subject variability, however, did not allow us to assess the specific effects of each type of stimulation on the original memory. Future experiments could conduct the same stimulation within one animal to assess the efficacy of certain forms of stimulation in eliciting artificial recall and its impact on the original memory and synaptic plasticity (Chen et al. 2019; Nabavi et al. 2014).

Overall, our results with regards to phase specific effects on engram activation on recall support the SPEAR model (Hasselmo et al. 2002), as well as a general role for theta oscillations in organizing memory recall in the hippocampus. The effectiveness of trough stimulating in eliciting recall is also consistent with previous experiments in rodents (Douchamps et al. 2013; Manns et al. 2007; Siegle and Wilson 2014) and human subjects (Kragel et al. 2020) performing memory tasks. Our results also support the general functional organization of the hippocampus and its relationship to theta oscillations. During the trough of theta (referenced to the pyramidal cell layer of CA1), a majority of the inputs to the CA1 region originate from CA3 (Mizuseki et al. 2009) and are responsible for the retrieval of stored memories. Inputs from CA3 converge at the *striatum radiatum* (SR) region of CA1 neurons (Brankačk and Buzsáki 1986; Li et al. 1994) and are accompanied by slow to mid-gamma range oscillation (30-85 Hz) (Schomburg et al. 2014). Activating engram neurons in DG at the trough of theta leads to in-phase activation of CA3 neurons and enhanced theta-gamma PAC (Figure 5). On the other hand, when stimulating at the peak of theta, inputs from EC (Mizuseki et al. 2009) terminating at the *lacunosum-moleculare* (LM) layer of CA1 (Amaral and Witter 1989) results in the encoding of new information. As a result, during peak activation of DG engrams, the out of phase activity that is conveyed through CA3 can compete with the stronger EC input (Basu and Siegelbaum 2015). These out of phase inputs to the LM and SR layers of CA1 can potentially disrupt the organization of CA1 activity (Figure 6). This interpretation is in line with the observation that the efficacy of peak stimulation is negatively correlated with theta-gamma synchrony (Figure 5D). Nevertheless, network measures of activity in CA1 and CA3 that incorporate direct measures of neuronal output are required to demonstrate the network effects of in phase and out of phase engram activation.

**Figure 6.**
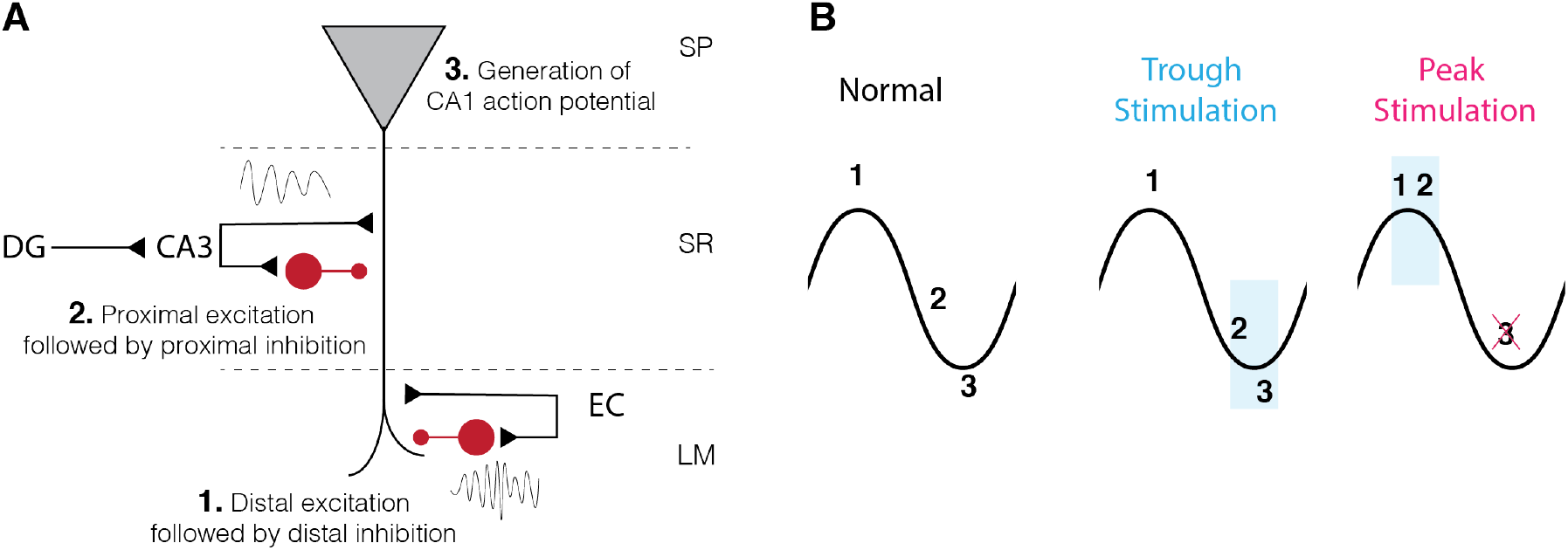
Phase-specific inputs to CA1 pyramidal neurons interact with the sequence of activity in different hippocampal networks. ***A***. Schematic showing CA3 and EC inputs to CA1 pyramidal cells and interneurons. CA3 input at the trough of theta is accompanied by slow-mid gamma oscillations, while EC input during the peak of theta oscillations is accompanied by higher frequency gamma oscillation. ***B***. During physiological activity in hippocampus, EC input, followed by CA3 input results in the activation of CA1 neurons. In the case of trough activation of engram neurons, the order is preserved, while in the case of the peak stimulation, the simultaneous activation of EC and CA3 inputs disrupts CA1 activity.

Results presented here open the door for an exciting line of research with regards to a role of theta oscillations in the context of memory processing. Future studies using phase-specific activation of engram neurons will greatly benefit from combining stimulation with calcium imaging (Grienberger and Konnerth 2012) or high density electrode arrays (Jun et al. 2017), which could provide single neuron- and network-based mechanisms for hippocampal theta oscillation function during memory processing. Another potential research route for probing the role of theta oscillations in memory gating is through a comparison of encoding and recall engram neurons in the CA3 and EC regions, respectively. For example, we expect tagged CA3 neurons, which are associated with the recall of a fearful memory, to be more robustly reactivated when stimulating at the trough of theta. To our knowledge, however, no studies to date have demonstrated the successful artificial reactivation of memories in CA3. We also hypothesize that EC inputs to the hippocampus can be tagged during the encoding of an experience, with subsequent peak stimulation of the tagged neurons during a second experience disrupting the encoding of that event.

In conclusion, our systematic investigation of engram neuron activation using different modes of stimulation provides new insights regarding the impact of stimulation frequency and phase on engram reactivation, as well as the utility of using a closed loop photo-stimulation approach.

## IV Methods

### Animals

All procedures were done in accordance with the National Institutes of Health Guide for Laboratory Animals and were approved by the Boston University Institutional Animal Care and Use and Biosafety Committees. We exclusively used adult C57bL/6 wild type male mice (aged 4-8 months). Exclusion of female mice from the study was based on the observation that female mice expressed elevated anxiety relative to male mice, which made assessing fear responses very difficult. Animals were acquired from The Jackson Laboratory.

### Sampling and exclusion criteria

No statistical methods were used to determine the sample size; the number subjects per group was determined based on previously published studies. We used 53 male mice in the current study, with 10 serving as control and 43 as experimental animals. Animals were randomly assigned to the experimental versus control group. Mice were included in the analysis based on pre-defined factors regarding viral expression, effects of the light stimulation and the quality of LFP recording. If the experimental animals did not show at least 5% increase in their freezing as a result engram reactivation on two out of four days of the experiment, they were excluded. The exclusion was further confirmed based on virus expression using post-hoc analysis of brain slice slides. Further, since it was crucial for phase-locked stimulation to have an LFP signal with low noise, if the quality of LFP was deemed unsatisfactory to drive reliable phase-locked stimulation, the animal was removed from further analysis. The exclusion left 17 experimental animals and 9 control animals for the analysis.

### Surgeries

To express the virus, as well as implant the fiber optics and the electrode, animals underwent stereotaxic surgery 3 to 4 weeks prior to the start of the behavioral experiments. Twenty-four hours prior to the surgery, animals were put on 40 mg/kg doxycycline diet. For the surgery, mice were anesthetized with isoflurane (1.8 to 2%) vaporized in room air. Bilateral holes were drilled above the dorsal DG at −2.2 mm anterioposterior (AP); ± 1.3 mm mediolateral (ML) from Bregma. To express channel rhodopsin in engram neurons, animals were bilaterally injected with AAV_9_-TRE-ChR2-EYFP acquired from the Massachusetts Institute of Technology, at a depth of 1.8 mm dorsoventral (DV) from the surface. A total of 200 nl of the virus was injected using a 10-nl syringe (World Precision Instruments (WPI)) fitted with a 33-gauge needle (NF33BL; World Precision Instruments), at a speed of 50 nl/min that was controlled via a microsyringe pump (UltraMicroPump 3–4; WPI). Post injection, one side was implanted with a fiber optic cannula (200 μm core diameter, 0.39 numerical aperture; Doric Lenses). Following that, the other side was implanted with a costume single LFP electrode (diameter 125 μm, acquired from invivo1) glued to a fiber optic. Cannulas were implanted at -1.6 mm DV above the injection site. The electrode was targeted to stratum pyramidale layer of CA1 region of the hippocampus. Location of the electrode was validated post-hoc using Prussian Blue staining to identify tip of the electrode and ensure correct phase of theta was measured. Craniotomy was secured with a layer of metabond followed by dental cement. Postop animals received an intraperitoneal injection of the analgesia Buprenorphin (0.2–0.5 mg/kg) which was continued for 48 h post-surgery every 8–12 h.

### Behavioral Experiments

After recovery, prior to behavioral experiments, mice were handled daily to habituate them to the transportation and researchers. For the first few days, handling was accompanied with a small treat. Behavioral experiments started after animals were acclimatized to the experimenter, Behavioral testing was conducted in a 30.5 × 24 × 21 cm^3^ conditioning chambers (Med Associates). During all the trials, both fiber optics and the LFP electrode were plugged in to normalize the effects of the distress between all trials.

Two contexts were designed for the experiment. The fearful context B contained the bare chamber with metal rods on the bottom, aluminum side walls and a 20 kHz, 40 dB noise source. In the neutral context A, walls and the floor were covered with striped papers along with an ambient white light. Wooden cage bedding was present on the floor and walls were infused with an orange scent. Animals’ behavior was monitored using a near infrared camera.

The experiment started with 4 days of habituation for the animals. During which animals were exposed to context A while both fiber optics and the LFP electrode were plugged in. The trial was designed to be the same as the experimental trial. On each day animals stayed in the chamber for four 3-minute epochs alternating between no stimulation and stimulation. The habituation days got the animal acclimatized to the experimental environment, with data from the last day used as a baseline for the rest of the experiment.

Mice were taken off doxycycline 48 hour prior to the tagging experiment. To tag engram neurons, animals were introduced to context B, and after 5 minutes of exploration, the experimental group received four 1.2 mA electrical foot shocks over the next five minutes. Control group animals did not receive any foot shocks. Subjects were put back on the doxycycline immediately after the experiment. In following day, animals were put back in context B, to assess recall of the fearful behavior.

Testing the different stimulation setups was done in context A over 4 days. At the conclusion of the fourth day, mice were re-exposed to context B on day 5, after which mice were being sacrificed and perfused for histological analysis. This last step was performed to assess the long-term effects of artificial reactivations on the original memory.

### Reactivation of engram neurons

To test the effectiveness of different stimulation setups, engram neurons for each animal were optogenetically activated in the context A over four days. The order of different stimulations was randomized to control for possible effects of repeated stimulation using an equal number of animals for each stimulation at each specific order. Engram reactivation on each day consisted of four 3 minute epochs: no stimulation, stimulation, no stimulation, and last epoch of stimulation.

### Electrophysiology and optogenetics stimulation

LFP signals were gathered via custom electrodes created by inVivo1. Signals were recorded using a head-stage connected to a Molecular Devices, Axon Instrument amplifier and digitizer. Digitized signal was recorded at 1 kHz. The closed-loop algorithm implemented in RTXI (Lin et al. 2010), filtered the signal at 2-10 Hz and delivered a TTL pulse to drive a DPSS Single Longitudinal Mode 473 nm laser (optoEngine LLC) at a predicted time for stimulation (peak or trough or fixed frequency).

### Behavioral Scoring

In order to remove experimenter’s bias from the analysis, an open source python package named ezTrack (Pennington et al. 2019) was used to score the freezing during each trial. Parameters for the analysis of motion were manually tuned and kept consistent between trials of the same day in the same animal. For the mice to be considered freezing, it required the animal to remain still for at least 6 seconds. To ensure accuracy of the algorithm, random blinded trials were hand scored and compared with the automated analysis. Since there was no significant difference between the two, the automated analysis was kept for all trials.

### Slice preparation and histology

After completion of experimental manipulations, animals were introduced to context B for the last recall. Ninety minutes after testing, animals were transcardially perfused with cold PBS and tissue fixed using 4% paraformaldehyde (PFA) in PBS with ferrocyanide dissolved in the solution (10%). Brains were extracted and after 24-hour storage in 4% PFA, 50-100 μm slices were prepared. For EYFP staining, slices were incubated at 4°C with PBS and 0.2% triton and normal goat serum solution for 1 hour, followed by an incubation period with a primary anti GFP chicken antibody 1:1000 (Invitrogen, catalogue # A-10262). After 48 hours of incubation, slices were washed and then incubated with a secondary antibody 1:500, Alexa 488 goat-anti chicken (Invitrogen catalogue # A-11039) for one hour. Slices were washed and mounted on a slide with Vectashield (Vector Laboratories H-1000-10) containing a DAPI stain (stain for cell bodies). Olympus FV3000 was used for confocal imaging of slices and a z stack of 10 μm thick slices were acquired.

### Statistical analysis

For each of the statistical analysis both visual inspection of the distribution histogram and Shapiro-Wilk test of normality was performed. Following the determination of normal distribution, Levene’s test for equality of variance was performed to determine appropriate variables in the test. In all the cases, data were normal and paired or independent t-test was applied to test for significance. In cases of multiple comparison, the Bonferroni correction of multiple comparisons was used. All the statistical tests were done in Python, using custom scripts taking advantage of SciPy (Virtanen et al. 2020) package.

### LFP Analysis

LFP analysis was performed through custom Python scripts. Analysis took advantage of Scipy and a Python implementation (https://github.com/dz256/Matlab2Python) of the Chronux toolbox (Bokil et al. 2010). For the spectral analysis, zero-mean signal was bandpass filtered from 1 to 100 Hz using a 4^th^ order Butterworth filter. A notch filter was applied to remove the 60 Hz noise (4^th^ order Butterworth band-stop filter, with a center frequency of 60 Hz). Spectral content was estimated using a multi-taper method (9 tapers) with a 5 second sliding window, and 1 second overlap.

To identify the actual phase of theta, and determine sensitivity and specificity of the algorithm, post hoc analysis was performed. Data was filtered using fourth order Butterworth filter (4-10 Hz bandpass) and a Hilbert-transform was performed to determine the phase of signal. If the stimulation was within quarter cycle of the peak (0 degree) or trough (180 degree), it was considered on target for respective stimulations, otherwise it was considered off target. Sensitivity was calculated by dividing the number of stimulations by total number of in phase extrema detected. Specificity was calculated by dividing number of out of phase extrema not stimulated by the total number of out of phase extrema.

### Phase Amplitude Coupling (PAC) Analysis

A metric adopted from a previous study (Tort et al. 2010), termed the modulation index (MI), was applied to identify the coupling between gamma amplitude and theta phase. This metric is defined through comparing the amplitude distribution through a theta cycle with a uniform distribution. The phase from the Hilbert transform of the LFP filtered at theta range (4-10 Hz) was used as the phase signal. Amplitude of Hilbert transform of the LFP filtered at gamma frequency (65-85 Hz) was used as the amplitude distribution. Data was binned in twenty, 18-degree bins, with the amplitudes normalized by the average amplitude of the signal. This normalized distribution (P) was compared to a uniform distribution (U) using Kullback-Leibler (KL) divergence. MI was calculated by dividing the KL divergence by logarithm of the number of bins (N).

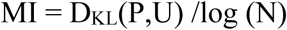

Comodulograms, like those in Figure 5, were created by calculating MI between theta and gamma by binning the phase frequencies in bands of 4 Hz, steps of 1 Hz and amplitude frequencies in bands of 20 Hz and steps of 5 Hz.

## V. Acknowledgements

We acknowledge the use Boston University Biomedical Engineering Department core micro and nano imaging facilities. We thank Dr. Michael Hasselmo, Dr. Christopher Harvey and Dr. David Boas for their feedback and guidance through the development of this research. This work was supported by NSF NRT: National Science Foundation Research Traineeship Program (NRT): Understanding the Brain (UtB): Neurophotonics DGE-1633516.

## VI. Data availability

Data collected for the purpose of this paper and the custom algorithms that were used in performing the analysis are available per request by contacting the corresponding author. Theta phase detection algorithm is accessible by contacting corresponding author, you can run the algorithm using the rTXI platform accessible through http://rtxi.org. Behavioral scoring was done using the ezTrack package available at github.com/DeniseCaiLab/ezTrack.

